# Impaired clock gene expression and abnormal diurnal regulation of hippocampal inhibitory transmission and spatial memory in a mouse model of Alzheimer’s disease

**DOI:** 10.1101/2021.02.12.430852

**Authors:** Allison R. Fusilier, Jennifer A. Davis, Jodi R. Paul, Stefani D. Yates, Laura J. McMeekin, Lacy K. Goode, Mugdha V. Mokashi, Thomas van Groen, Rita M. Cowell, Lori L. McMahon, Erik D. Roberson, Karen L. Gamble

**Author notes:** Corresponding Author: Karen L. Gamble, Department of Psychiatry and Behavioral Neurobiology University of Alabama at Birmingham, 1720 7th Ave S. Birmingham, AL 35294, w. These authors contributed equally: AR Fusilier and JA Davis.

## Abstract

Patients with Alzheimer’s disease (AD) often have fragmentation of sleep/wake cycles and disrupted 24-h (circadian) activity. Despite this, little work has investigated the potential underlying day/night disruptions in cognition and neuronal physiology in the hippocampus. The molecular clock, an intrinsic transcription-translation feedback loop that regulates circadian behavior, may also regulate hippocampal neurophysiological activity. We hypothesized that disrupted diurnal variation in clock gene expression in the hippocampus corresponds with loss of normal day/night differences in membrane excitability, synaptic physiology, and cognition. We previously reported that the Tg-SwDI mouse model of AD has disrupted circadian locomotor rhythms and neurophysiological output of the suprachiasmatic nucleus (the primary circadian clock). Here, we report that Tg-SwDI mice failed to show day-night differences in a spatial working memory task, unlike wild-type controls that exhibited enhanced spatial working memory at night. Moreover, Tg-SwDI mice had lower levels of Per2, one of the core components of the molecular clock, at both mRNA and protein levels when compared to age-matched controls. Interestingly, we discovered neurophysiological impairments in area CA1 of the Tg-SwDI hippocampus. In controls, spontaneous inhibitory post-synaptic currents (sIPSCs) in pyramidal cells showed greater amplitude and lower inter-event interval during the day than the night. However, the normal day/night differences in sIPSCs were absent (amplitude) or reversed (inter-event interval) in pyramidal cells from Tg-SwDI mice. In control mice, current injection into CA1 pyramidal cells produced more firing during the night than during the day, but no day/night difference in excitability was observed in Tg-SwDI mice. The normal day/night difference in excitability in controls was blocked by GABA receptor inhibition. Together, these results demonstrate that the normal diurnal regulation of inhibitory transmission in the hippocampus is diminished in a mouse model of AD, leading to decreased daytime inhibition onto hippocampal CA1 pyramidal cells. Uncovering disrupted day/night differences in circadian gene regulation, hippocampal physiology, and memory in AD mouse models may provide insight into possible chronotherapeutic strategies to ameliorate Alzheimer’s disease symptoms or delay pathological onset.

## INTRODUCTION

Alzheimer’s disease (AD), the most prevalent cause of dementia worldwide (Plassman et al., 2007), is characterized by development of pathological amyloid beta plaques and tau tangles throughout the brain, and regions associated with memory such as the hippocampus are particularly vulnerable. Patients with AD also exhibit subclinical epileptiform activity or seizures, indicative of network hyperexcitability (Palop and Mucke, 2016). This network hyperexcitability is seen in early, even preclinical, stages of AD (defined by normal cognition with positive biomarkers for amyloid pathology). In general, epileptiform activity is greater and seizure thresholds are lower during sleep/rest, resulting in increased frequency of seizures during that time of day (Gerstner et al., 2014; Spencer et al., 2016). Thus, network hyperexcitability may be exacerbated in AD patients because of documented disruptions in sleep-wake cycles, misaligned body temperature rhythms, and altered patterns of activity, all of which are driven by the circadian clock (Musiek et al., 2018; Volicer et al., 2001).

Circadian rhythms are endogenous, ∼24-h oscillations in physiology and behavior that are driven by the molecular clock. The molecular clock is a cellular transcription-translation feedback loop, involving the proteins BMAL1, PER1/2, and CRY1/2, present in nearly all cells of the body (Partch et al., 2014). The principal circadian pacemaker located in the hypothalamus, the suprachiasmatic nucleus (SCN), is responsible for coordinating rhythms throughout the body and in different regions of the brain (Schwartz et al., 1987 {Paul, 2019 #1128)}. In addition to physiological rhythms, cognitive performance also fluctuates across the course of the day, peaking shortly after waking and declining after several hours (Wright et al., 2006). Environmental disruptions, such as fragmented sleep or altered light cycles, and genetic manipulations to components of the molecular clock impair cognition (Knutsson, 2003; Snider and Obrietan, 2018; Wright et al., 2006; Wu et al., 2015; Wyatt et al., 1999). Similar to rhythms of behavior and cognition, electrophysiological properties fluctuate across the course of the day in many brain regions, including the hippocampus (Naseri Kouzehgarani et al., 2019; Paul et al., 2019). Rodent studies have shown that hippocampal long-term potentiation (LTP), a synaptic correlate of learning and memory (Whitlock et al., 2006), is enhanced during the active period compared to the inactive period (Besing et al., 2017; Chaudhury et al., 2005). In fact, whole-body deletion of *Bmal1* impairs LTP (Wardlaw et al., 2014). At the cellular level, rat CA1 pyramidal cell resting membrane potential is more depolarized at night relative to day (Naseri Kouzehgarani et al., 2019). Clock-controlled genes expressed in the hippocampus may underlie this link given that over 600 transcripts show circadian variation in the hippocampus, including those encoding ion channels and synaptic proteins such as *Kcna4, Clca2, Trpc7, Kcnk3, Gria3*, and *Nlgn1* (Renaud et al., 2015).

We recently found circadian impairment in the central SCN clock in the Tg-SwDI model of AD (Paul et al., 2018) – these mice express the human *APP* gene with the Swedish (K670N/M671L), Dutch (E693Q), and Iowa (D694N) mutations at levels similar to the levels of mouse APP (Davis et al., 2004). We showed that these mice do not have hyperactive locomotor activity frequently observed in other mouse models of AD (Sheehan and Musiek, 2020) but do show a significantly shorter free-running period than age-matched controls at 3, 6, and 10 months of age. In addition, Tg-SwDI mice at 6-8 months of age have greatly reduced day-night differences in SCN action potential firing rate (compared to control mice), primarily due to a lower daytime firing rate (Paul et al., 2018). Here, we asked whether these circadian changes extend to other brain regions, particularly the hippocampus. Tg-SwDI mice develop fibrillar amyloid deposition specifically in the cerebral microvasculature rather than in the parenchyma, and this pattern of deposition is associated with impaired spatial learning and memory (Davis et al., 2004; Xu et al., 2007; Xu et al., 2014). To our knowledge, examination of day/night differences in hippocampal function or physiology has yet to be assessed in this model or other models of AD. Therefore, in the present study we sought to determine whether altered expression of key molecular clock components in the hippocampus corresponds with loss of day/night differences in membrane excitability, synaptic physiology, and cognition.

## MATERIALS AND METHODS

### Animals

All animal procedures followed the Guide for the Care and Use of Laboratory Animals, U.S. Public Health Service, and were approved by the University of Alabama at Birmingham Institutional Animal Care and Use Committee. Mice of both sexes were maintained on a 12:12 light/dark cycle with ad libitum access to food (LabDiet Rodent 5001 by Purina) and water. Mice expressing the human amyloid precursor protein with the familial Swedish (K670N/M671L), Dutch (E693Q), Iowa (D694N) homozygous mutations (Tg-SwDI; Davis et al. 2004) were bred at the University of Alabama at Birmingham (UAB) on a congenic on a C57BL/6 background. Mice were compared to age matched C57BL/6 wild type (WT) mice from the National Institute on Aging Aged Rodent Colony or from the C57BL/6 colony at UAB. Control, C57BL/6 mice used in the electrophysiology experiment in Figure 4 also carried the *Per1*::GFP transgene (Paul et al., 2016).

### Spatial working memory

Spatial working memory was tested during the day (Zeitgeber time (ZT) 2) and night (ZT 14) with a T-maze, spontaneous alternation protocol (adapted from (Deacon and Rawlins, 2006)). Mice were handled for 4 consecutive days prior to testing to reduce anxiety from being handled during the test. Mice were placed in the maze at the base of the T arm and allowed to freely choose an arm of the T. Upon entry to an arm, a gate was closed, trapping the mouse in the arm for 30 seconds. The mouse was removed, the gate opened, and the mouse returned to the base of the maze. An alternation was recorded if the mouse entered the opposite arm as the last entry. Each mouse performed 5 trials, and an alternation percentage was calculated as (# alternations / # of trials). Males and females were tested separately, and WT and AD were interleaved during trials. During night testing, dim red light (< 5.0 lux) was used for visualization. Animals were only excluded from analysis if there was significant cage disruption during the time period of the test (i.e., excessive fighting in the animal cage) or if there was a notable directional bias in arm choices (i.e. animals only used the left arm in all 10 entries).

### Electrophysiology

*Voltage clamp recordings* at 4 months of age, mice were euthanized with cervical dislocation and rapid decapitation, brains were removed, and 350 μm coronal slices were prepared using a VT1000P vibratome (Leica Biosystems). Slices were cut in ice cold cutting solution containing the following (in mM): 110 choline chloride, 25 glucose, 7 MgCl2, 2.5 KCl, 1.25 Na2PO4, 0.5 CaCl2, 1.3 Na-ascorbate, 3 Na-pyruvate, and 25 NaHCO3, bubbled with 95% O2/5% CO2. Slices were rested at room temperature for at least one hour in recovery solution containing the following (in mM): 125 NaCl, 2.5 KCl, 1.25 Na2PO4, 2 CaCl2, 1 MgCl2, 25 NaHCO3, 25 glucose and 2 kynurenic acid bubbled with 95% O2/5% CO2. Recordings were made in standard artificial cerebral spinal fluid (ACSF) containing the following (in mM): 125 NaCl, 2.5 KCl, 1.25 Na2PO4, 2 CaCl2, 1 MgCl2, 25 NaHCO3, and 25 glucose bubbled with 95% O2/5% CO2. Patch pipette solution contained (in mM): 140.0 CsCl, 10.0 EGTA, 5.0 MgCl2, 2.0 Na-ATP, 0.3 Na-GTP, 10.0 HEPES, and 0.2% biocytin (pH 7.3, 290 mOsm), and QX-314 (sodium channel antagonist, 50 μM) added at time of use. Examples of biocytin filled pyramidal neurons from both WT and Tg-SwDI mice are shown in Supplementary Figure 1. Patch pipettes (BF150-086; Sutter Instruments, Novato, CA) were pulled on a Sutter P-97 horizontal puller (Sutter Instruments, Novato, CA) to a resistance between 3-5 MΩ. Recordings were performed in a submersion chamber with continuous perfusion of oxygenated ACSF at 2.5 mL/min. The blind patch technique was used to acquire whole-cell recordings from CA1 pyramidal neurons. Neuronal activity was recorded using an Axopatch 200B amplifier and pClamp10 acquisition software (Molecular Devices, Sunnyvale, CA). Signals were filtered at 5 kHz and digitized at 10 kHz (Digidata 1440). Spontaneous inhibitory post-synaptic currents (sIPSCs) were recorded between ZT 4-10 or ZT 13-19 and pharmacologically isolated with bath perfusion of NBQX (AMPAR antagonist, 10 μM, Hello Bio) and CPP (NDMAR antagonist, 5 μM, Hello Bio). All cells were dialyzed for 5 min prior to experimental recordings and held at −70 mV. Cells used for analysis had access resistance less than 30 MOhms that did not change by more than 20% for the duration of recording.

#### Current Clamp Recordings

At 6-7 months of age, mice were euthanized via cervical dislocation at ZT 0-1 for day or 11-12 for night recordings and coronal hippocampal slices (300 µm) were prepared using a Campden 7000SMZ (World Precision Instruments, Lafayette, IN). Slices were cut in cold cutting solution containing the following (in mM): 250 sucrose, 26 NaHCO3, 1.25 Na2HPO4-7H2O, 1.2 MgSO4-7H2O, 10 glucose, 2.5 MgCl2, 3.5 KCl bubbled with 95% O2/5% CO2. Slices were transferred to a beaker containing 50% sucrose saline and 50% normal saline (in mM: 130 NaCl, 20 NaHCO3, 1 Na2HPO4-7H2O, 1.3 MgSO4-7H2O, 10 Glucose, 3.5 KCl, 2.5 CaCl2) at room temperature for 20 min and then transferred to an open recording chamber (Warner Instruments, Hambden, CT) that was continuously perfused at a rate of 2.0 ml/min with normal saline, bubbled with 5% CO2 / 95% O2 and heated to 34 ± 0.5 °C. Patch pipette solution contained (in mM): 135 K-gluconate, 10 KCl, 10 HEPES, 0.5 EGTA; pH 7.4. Patch pipettes (BF150-110-10; Sutter Instruments, Novato, CA) were pulled on a Sutter P-97 horizontal puller (Sutter Instruments, Novato, CA) to a resistance between 4-6 MΩ. Neuronal activity was recorded using an Multiclamp 700B amplifier and pClamp10 acquisition software (Molecular Devices, Sunnyvale, CA). Signals were filtered at 10 kHz and digitized at 20 kHz (Digidata 1440A). Whole-cell current clamp recordings were made from CA1 pyramidal cells using blind patch technique between ZT 2-8 or ZT 13-19 for day and night recordings, respectively. Resting membrane potential was measured in gap free current clamp mode (as in (Paul et al., 2016). Neuronal excitability was studied by injecting progressive steps of depolarizing current from rest (0 pA to 280 pA at 20 pA increments) and counting the number of action potentials firing during each 800ms current step. For recordings in *Per1::GFP* mice, slices were treated with GABAR antagonists (10µM bicuculline and 1µM CGP55848) or vehicle (0.002% DMSO) continuously in the bath for the duration of the recording.

### Quantitative real-time PCR (qRT-PCR)

Quantitative real-time PCR was performed as previously described (Lucas et al. 2012). At 4 months of age, mice were euthanized via cervical dislocation at ZT 2, 8, 14, and 20 with the aid of night vision goggles for enucleation at dark time points. Brains were removed and hippocampus was extracted and stored at −80°C. Tissue was homogenized in TRIzol (Thermo Fisher Scientific) using the Omni Bead Ruptor Homogenizer (OMNI International, Kennesaw, GA, USA). RNA was then isolated using the choloform-isopropanol method following the manufacturer’s instructions (Invitrogen). RNA concentration and purity were assessed using a NanoDrop One (Thermo Fisher Scientific). Equivalent amounts of RNA (1 µg) were treated with DNase I (Promega) at 37°C for 30 minutes, DNase Stop solution at 65°C for 15 minutes and then RNA was reverse-transcribed using the High-Capacity cDNA Reverse Transcription Kit (Thermo Fisher) at 37°C for two hours. Samples were then stored at −20°C. Transcripts were measured using mouse-specific primers from Applied Biosystems and JumpStart Taq Readymix (Sigma, St. Louis, MO, USA) using a protocol with an initial ramp for 10 min at 95°C and 40 subsequent cycles of 95°C for 15 sec and 60°C for one min. Relative concentration of transcript was calculated in comparison to a standard curve generated from pooled cDNA samples and then diluted (1:5, 1:10, 1:20, 1:40; calibrator method). These values were then normalized to 18S (Hs99999901_s1) and expressed as mean +/- SEM. 18S values were analyzed to ensure no difference across time of day or genotype prior to normalization. The Applied Biosystems primer/probe sets used are as follows: beta-actin (Mm00607939_s1), Per1 (Mm00501813_m1), Per2 (Mm00478099_m1), Rev-erbα (Mm00520708_m1), Cry1 (Mm00514392_m1), Cry2 (Mm01331539_m1), Bmal1 (Mm00500223_m1), Bmal2 (Mm00549497_m1).

### Immunohistochemistry (IHC) and western blots

*IHC* At 4 months of age, mice were anesthetized with isoflurane and transcardially perfused with ice-cold PBS between ZT 3 and ZT 6. The whole brain was extracted and fixed in 4% paraformaldehyde for ∼24 h at 4°C, and then cryoprotected in 15% sucrose in PBS for ∼24 h followed by 30% sucrose in PBS for another 24 h at 4°C. Fixed, cryoprotected brains were then frozen on dry ice and stored at −80°C until 40-µm serial coronal slices were sectioned with a cryostat. For antibody labeling, sections were washed in PBS at RT for 3 × 10 min before blocking for 30 min in a PBS solution containing 5% Normal Donkey Serum (NDS) and 0.25% Triton X-100, followed by overnight incubation at 4°C with primary antibodies against GAD-67 (Millipore, 1:2000) and parvalbumin (Swant, 1:2000) in PBS containing 1% NDS and 0.1% Triton X-100. Sections were washed in 0.1% Triton X-100 PBS for 3 × 10 min before incubation with either donkey anti-rabbit Alexa fluor 488 (Invitrogen, 1:1000) or donkey anti-mouse Alexa fluor 555 (Invitrogen, 1:1000) secondary antibody in PBS containing 1% NDS and 0.1% Triton X-100. Sections were then washed in PBS and mounted on glass slides and coverslips with ProLong Gold Antifade Mounting media containing DAPI. Slides were imaged with a Nikon A1R confocal microscope using 10X and 20X objectives. Images were analyzed using ImageJ. For GAD analysis, background noise was automatically subtracted in ImageJ and three, identical rectangular regions of interest (ROI) were drawn in stratum oriens, stratum pyramidale, and stratum moleculare of area CA1 in dorsal hippocampus. The mean intensity of each ROI was measured in the channel of interest. For PV image analysis, PV^+^ cell counts were made in ImageJ using the multi-point tool.

*Western Blots* At 4 and 9 months of age, mice were euthanized via cervical dislocation at ZT 2, 8, 14, and 20 with the aid of night vision goggles for enucleation at dark time points. Brains were removed and whole hippocampus was extracted. Hippocampi were placed in ice-cold sucrose homogenization buffer (5 mM Tris-HCl (pH 7.4), 0.32 M sucrose, with a protease inhibitor tablet (Complete Mini, Roche Diagnostics, Mannheim, Germany)), minced using a scalpel blade, snap frozen in liquid nitrogen, and stored at −80°C until ready for homogenization. Samples were homogenized in ice-sold sucrose homogenization buffer using a Power Gen 125 homogenizer (Thermo Fisher Scientific, Waltham, MA). Protein concentrations were determined using a bicinchoninic acid (BCA) assay kit (Thermo Fisher Scientific, Pierce™ BCA Protein Assay Kit) and 15 µg homogenate were loaded into 4-12% Bis-tris gels (Invitrogen). Gels were transferred onto nitrocellulose immunoblotting paper using Bio-Rad Trans-Blot Turbo transfer system at 1.3 Amp and 25 V for 12 min. Blots were blocked in blocking buffer for 1 hour at room temperature and probed overnight at 4°C for PER2 (1:1000 in 5% BSA and TBST; Alpha Diagnostic Intl., Inc., #PER21-A), BMAL1 (1:1000 in 5% BSA and TBST, Signalway Antibody, LLC, #21415,), GSK3β (3D10) (1:1000 in 5% milk and TBST, Cell Signaling Technology, #9832), phospho-GSK3β (Ser9) (1:1000 in 5% BSA and TBST, Cell Signaling Technology, #93360), and GAPDH (1:5000, Cell Signaling, #2118). The appropriate IRDye secondary (Li-Cor Biosciences) was applied, and then blots were imaged and analyzed using Li-Cor Image Studio Lite version 5.2. Comparisons across immunoblots were achieved using an average of two or three inter-blot control samples on each blot. For every blot, all bands were normalized to the inter-blot average then to the loading control (GAPDH or GSK3β).

### Statistical Analysis

Statistics were calculated using SPSS software (version 25) and GraphPad prism 6 software. Assumptions of parametric tests, including normality and homogeneity of variance were determined, and if violated, nonparametric or ordinal tests were done. Ordinal regression was used for behavioral data analysis, stratified by sex (see behavioral methods above), with post hoc comparisons and Holmes corrected alpha. RNA and protein expression data were first assessed for a main effect of sex, and if significant, two-way ANOVAs were performed on data stratified by sex. In the absence of sex differences, two-way ANOVAs were performed with both sexes pooled. All significant interactions were followed by Tukey’spost hoc comparison. Current clamp data were analyzed with three-way ANOVAs with repeated measures followed by planned comparisons of each current step. Voltage clamp data were analyzed with Kolmogorov-Smirnov tests with Bonferroni correction, and histology data were compared with independent samples t-tests. Unless otherwise specified, significance was ascribed at p < 0.05. The data that support the findings of this study are available from the corresponding author.

## RESULTS

### Tg-SwDI mice lack normal diurnal patterns in spatial working memory

As previously mentioned, cognitive performance fluctuates across the day-night cycle, typically peaking during the time an animal is awake (Wright et al., 2012). As rodents are nocturnal, cognitive performance is better during the night than during the day (Shimizu et al., 2016; Snider et al., 2018), a phenomenon which persists even if mice are housed in constant darkness (Snider et al., 2016). Thus, testing at both times of day is important when assessing cognitive function. Tg-SwDI mice have spatial memory deficits (i.e., Barnes maze performance) when vascular amyloid deposits are first detected in the subiculum (∼3 months of age; (Xu et al., 2007; Xu et al., 2014)), although deficits in other cognitive tasks such as novel object recognition are not detected even at older ages (Robison et al., 2020; Robison et al., 2019). Importantly, day/night differences have yet to be assessed in Tg-SwDI mice on any memory task at any age. To address this question, we examined spontaneous alternation in a T-maze, a task with demonstrated day/night differences in mice (Davis et al., 2020) and other rodents (Fernandez et al., 2014; Ruby et al., 2015). This task employs trials, thus avoiding the confound of differences in the number of arm entries (Robison et al., 2019), and does not require multiple days of training that could disrupt sleep and circadian rhythms. Male and female Tg-SwDI mice (4 months and 9 months) were compared to control mice (day vs night) in four separate experiments. Day/night differences in alternation scores for female, 4-month-old mice depended upon genotype (Figure 1A and B, Supplementary Table 1; two-way ANOVA, interaction, p < 0.05). Control mice had significantly higher alternation at night compared to day (p < 0.05), but Tg-SwDI mice had no day/night difference due to significantly lower alternation at night compared to alternation of WT mice at night (p < 0.05). Males at 4 months of age showed better performance at night than during the day (Figure 1B main effect, p < 0.05); however, the interaction with genotype failed to reach significance (p = 0.20). At 9 months, day-night differences for both the male and female mice depended upon genotype (Figure 1C and D, p < 0.05). Although male and female control mice exhibited higher alternation at night (p < 0.05), neither male nor female Tg-SwDI animals showed a significant day/night difference in alternation (p > 0.38). These results show that, in general, diurnal rhythms in spatial working memory are impaired in this AD model.

**Figure 1.**
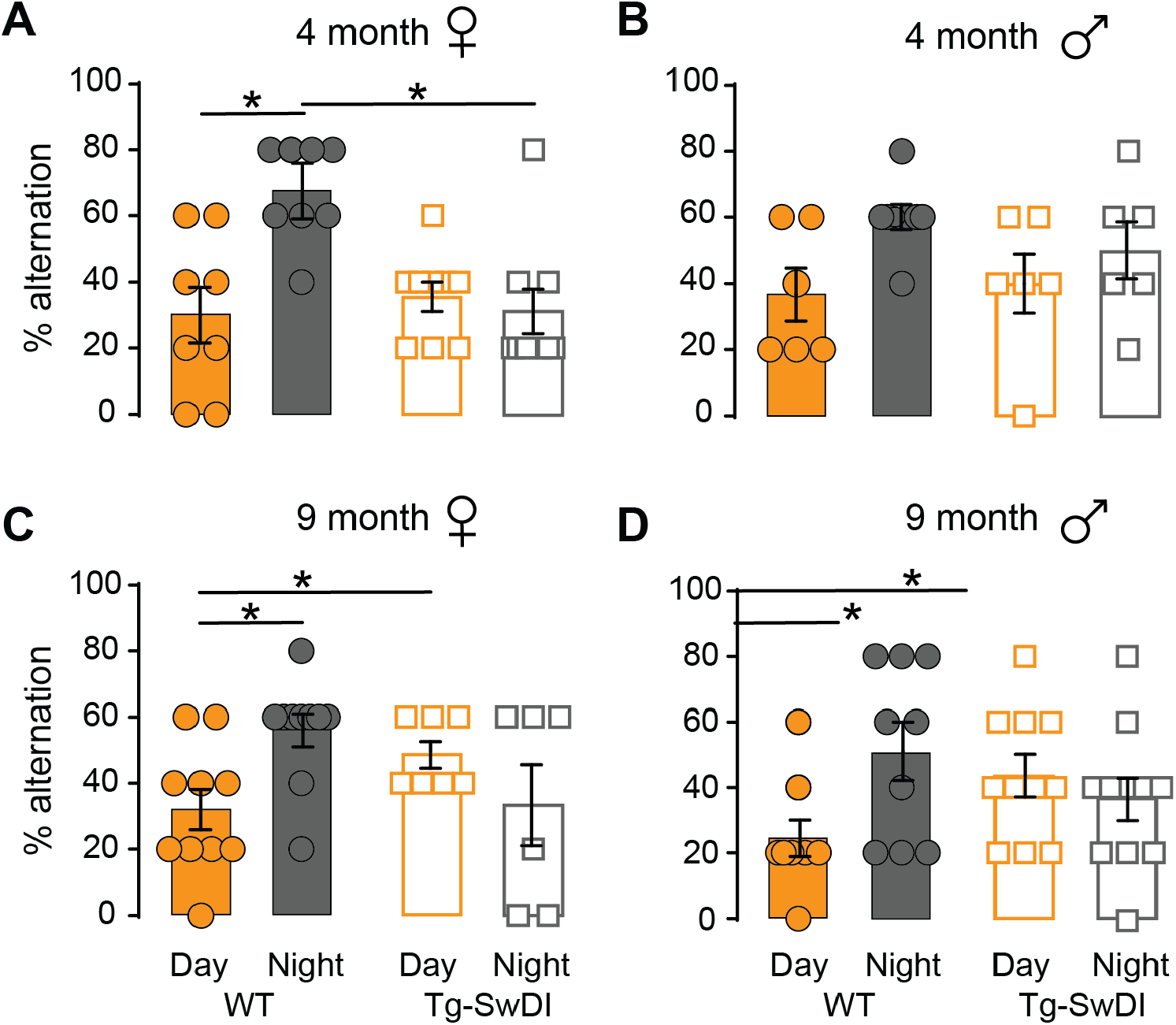
Loss of day/night differences in spatial working memory in 4-month and 9-month AD mice. Percent alternation at ZT 2 vs ZT 14 in WT and AD mice. 4-month WT and Tg-SwDI female (A) and male (B) mice tested during day (orange) and night (grey). WT mice had significant day/night difference in percent alternation (p < 0.05), and at night, Tg-SwDI mice had lower alternation percentages than WT mice (p < 0.01). 9-month WT and Tg-SwDI female (C) and male (D) mice tested during day (orange) and night (grey). Post-hoc analysis indicated a loss of day/night differences in both male and female Tg-SwDI mice (p = 0.386 and 0.521, respectively) and slightly higher alternation of older Tg-SwDI mice (males and females) compared to WT during the day (p < 0.02) and a tendency for lower alternation compared to WT females during the night (p = 0.076). For A, B, C, and D, a mixed model ordinal regression indicated significant main effects for time of day for all 4-month mice (p < 0.05), and significant Time of Day X Genotype interactions for 4-month females and all 9-month mice (p < 0.05, n = 34 (4-month females), 26 (4-month males), 33 (9-month females), 40 (9-month males)). See Supplementary Table 1 for statistical results of the regression.

### Alterations to the hippocampal molecular clock in Tg-SwDI mice

A functional circadian system is important for learning and memory (Ruby et al., 2008; Snider et al., 2016), and the Tg-SwDI AD model shows impaired spatial learning and memory (Figure 1; (Xu et al., 2007)) along with disrupted circadian wheel running behavior and altered SCN activity (Paul et al., 2018). Thus, we sought to determine whether Tg-SwDI mice also have disruptions of the hippocampal molecular clock. To test this hypothesis, we extracted whole hippocampus from 4-month Tg-SwDI males and females and age-matched WT controls at ZT 2, 8, 14, and 20, and performed qRT-PCR to examine mRNA expression of *Bmal1, Bmal2, Cry1, Cry2, Per1, Per2*, and *Rev-erbα*. All data for gene transcripts and proteins were stratified or pooled by sex based on a significant or non-significant (respectively) main effect of sex. All results of the two-way ANOVAs (Genotype X Time-of-Day) are reported in Supplementary Table 2.

There were no sex differences in mRNA expression of *Bmal1, Bmal2, Cry1*, and *Per1*, and thus, samples from males and females were pooled. Tg-SwDI mice had overall reduced expression of these genes compared to controls (Genotype main effect, p < 0.05; Figure 2A-C and E). Although diurnal variation was not evident for expression of these genes, *Cry1* expression appeared to show variation over time of day, but this effect failed to reach statistical significance (main effect of Time of Day, p = 0.089). Gene expression of *Cry2, Per2, and Rev-erbα* was stratified by sex for analysis due to a significant main effect of sex. Compared to control females, *Cry2* mRNA expression from Tg-SwDI females was generally increased (p < 0.05, Figure 2D), and *Per2* and *Rev-erbα* expression was decreased (p < 0.05, Figure 2F, and G). Expression of *Per2* showed diurnal variation in both females (Figure 2F) and males (Supplementary Figure 2A) with expression higher at both night time points (ZT 14 and ZT 20) compared to one or both of the day time points (females, ZT 2 and 8 vs ZT 14 and 20, p < 0.05; males, ZT 2 vs ZT 14 and 20, p < 0.05). Finally, *Rev-erbα* mRNA expression from male Tg-SwDI mice were significantly lower (regardless of time) than that of control mice (p < 0.05, Figure 2H). Expression of the control gene, 18s, did not vary across time of day (Supplementary Table 2).

**Figure 2.**
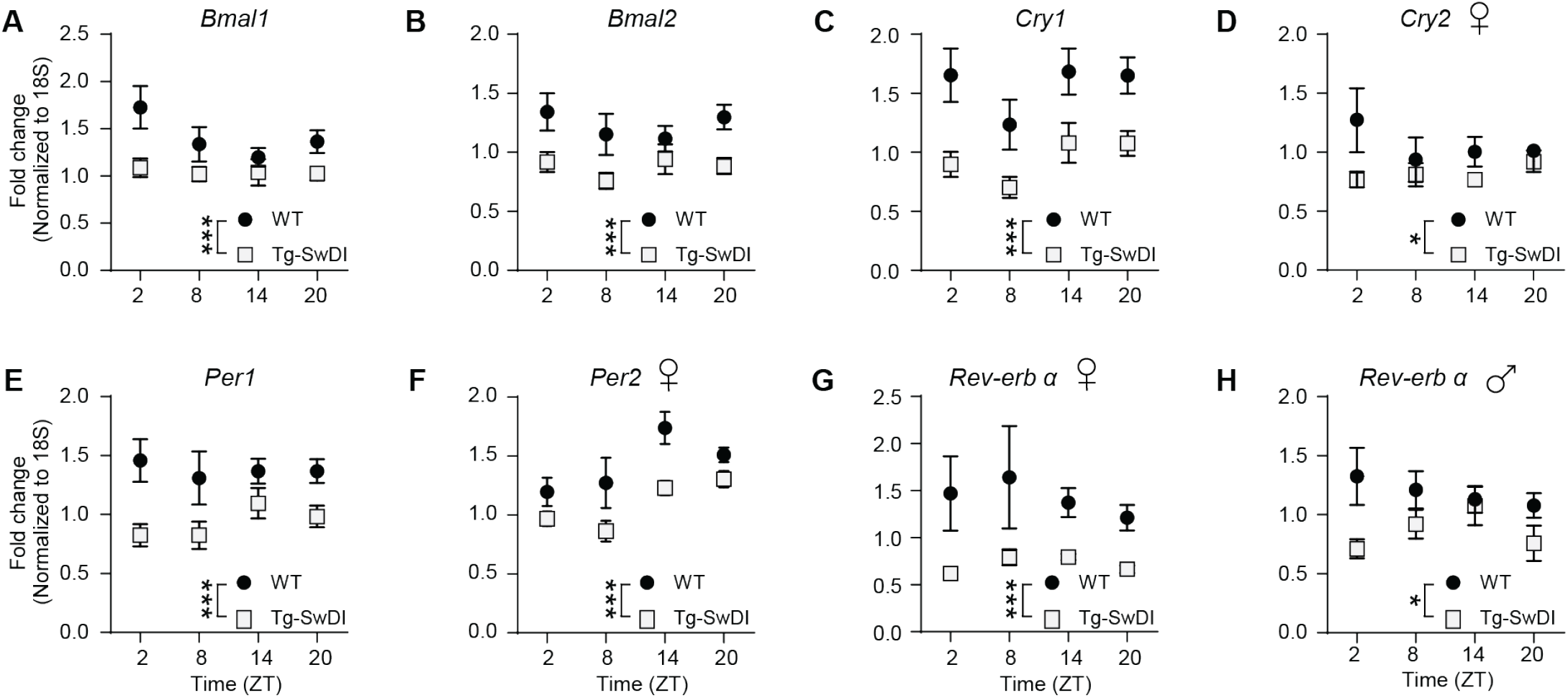
Reduced circadian gene mRNA expression over the day/night cycle in 4-month AD mice. Transcript levels from WT (filled circles) and Tg-SwDI (open squares) mice at ZT 2, 8, 14, and 20 for *Bmal1* (A), *Bmal2* (B), *Cry1* (C), *Cry2* females (D), *Per1* (E), *Per2* females (F), *Rev-erbα* females (G) and males (H). Two-way ANOVA indicated significant main effects of Genotype for *Bmal1, Cry1*, and *Cry 2* females, *Per1, Per2* females, and *Rev-erbα*. A significant Time of Day main effect was observed for *Per2* females (p < 0.05). No significant Time of Day X Genotype interactions were observed. Males and females were combined unless indicated (n = 5-10 mice per Sex/Time/Genotype group). See Supplementary Table 2 for statistical results.

To test if the changes in circadian clock gene mRNA expression also manifested at the protein level, we assessed hippocampal protein levels of two core clock components, PER2 and BMAL1, at both 4 months and 9 months. Given no significant effect of Sex, data from males and females were pooled for two-way ANOVAs (Supplementary Table 3). At 4 months, PER2 levels were significantly reduced in Tg-SwDI mice compared to controls (p < 0.05, Figure 3A). Regardless of genotype, PER2 levels showed diurnal variation with higher expression at ZT 20 than ZT 8 (Tukey, p < 0.05). For BMAL1, diurnal variation was dependent upon genotype, such that at ZT20, BMAL1 levels were higher in Tg-SwDI mice than in control mice (interaction, p < 0.05; Tukey’s HSD, p < 0.001; Figure 3B). At ZT14, there was a trend for BMAL1 levels from Tg-SwDi to be lower than WT (p = 0.57). By 9 months, overall levels of both PER2 and BMAL1 were significantly lower compared to control samples (main effect, p < 0.05; Figure 3C-D). At this age, there was a trend for diurnal variation in PER2 and BMAL1 levels (p = 0.079 and p = 0.088, respectively.

**Figure 3.**
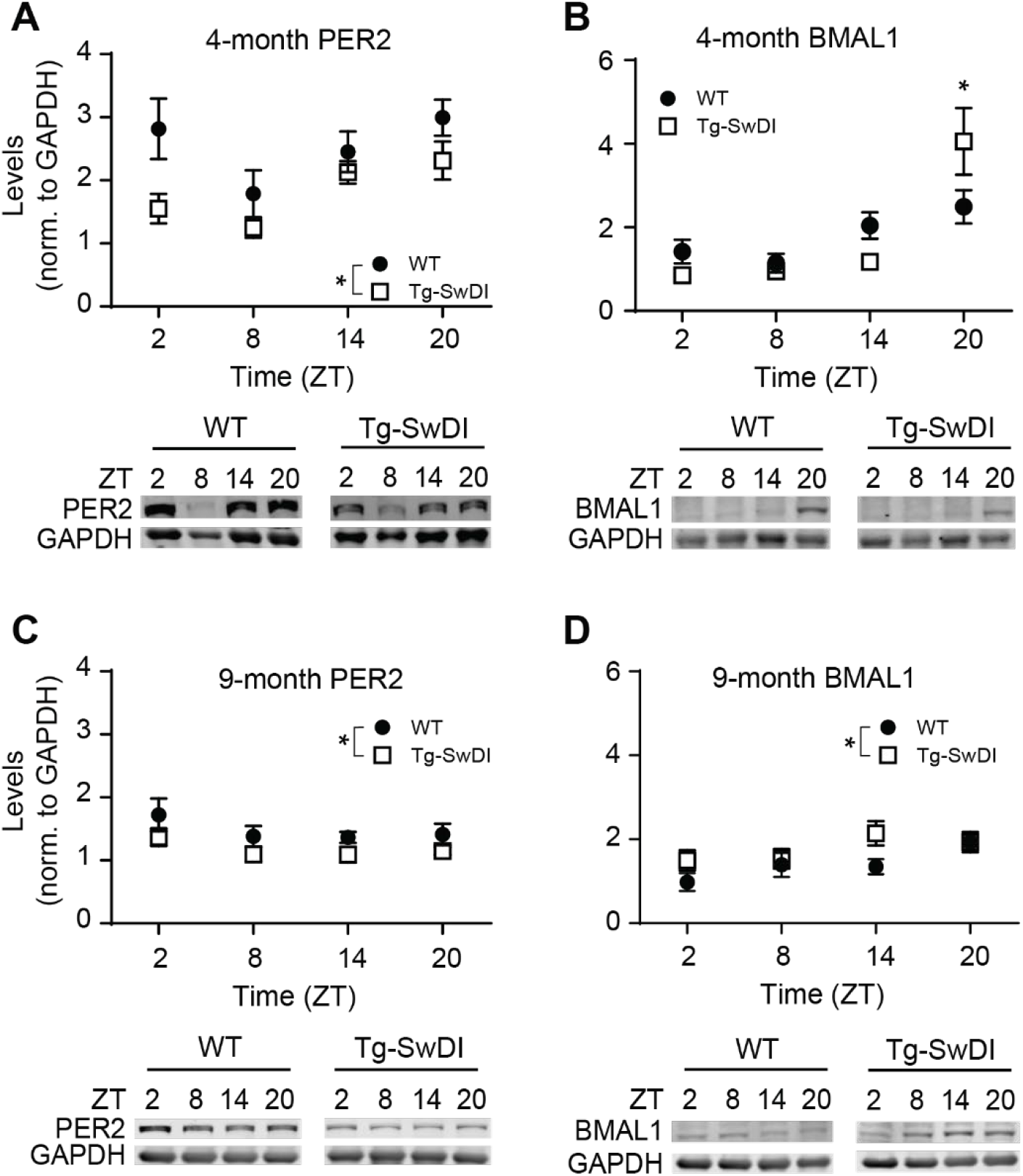
Loss of day/night variation in clock protein expression in Tg-SwDI mice. Normalized expression for PER2 and BMAL1 in 4-month (A and B) and 9-month (C and D) Tg-SwDI mice (open squares) and age-matched WT controls (solid circles). For 4-month mice, two-way ANOVA indicated significant main effects of Genotype and Time of Day for PER2 (A; p < 0.05) but no interaction (n = 9-15 mice/group). Levels of BMAL1 (B) varied by both Time-of-Day and Genotype (interaction, p < 0.01) with higher levels in AD mice than WT mice at ZT 20 (p < 0.05) and a trend for AD mice to have lower levels than WT mice at ZT 14 (p 0.057). Both Genotypes showed highest expression at ZT 20 (WT: ZT 20 vs ZT 8, p < 0.05; AD, ZT 20 vs all other ZTs, p < 0.05; n = 9-16 mice/group). For 9-month mice, two-way ANOVA indicated a significant main effect of Genotype for PER2 (C; p < 0.05) and a trend for Time of Day (p = 0.08), and no interaction (n = 10-16 mice/group). For BMAL1 (D), a significant main effect of Genotype (p < 0.05) revealed higher levels in AD mice than WT mice with a trend for a Time-of-Day effect (p = 0.088) but no significant interaction (p = 0.523). See Supplementary Table 3 for statistical results.

Phosphorylation of glycogen synthase kinase 3 beta (GSK3β), a second messenger protein important for spatial memory (Hernandez et al., 2002), is regulated in a circadian manner in the hippocampus (Besing et al., 2017). Moreover, GSK3β phosphorylates various components of the molecular clock (Besing et al., 2015; Iitaka et al., 2005; Iwahana et al., 2004). Beyond involvement in both circadian rhythms and memory, phosphorylated (inactive) GSK3β is reduced in AD (Fuentealba et al., 2004). At 4 months, phosphorylation of GSK3β (Supplementary Figure 3A) peaked at ZT 8 in both genotypes, although no Genotype difference was observed at this age (data for males and female pooled given no effect of Sex; Time-of-Day main effect, p < 0.05; Genotype main effect, p = 0.14; Supplementary Table 3). At 9 months, we detected no diurnal variation in GSK3β phosphorylation (data stratified by Sex, for both males and females, p > 0.19, Supplementary Figure 3B and C). Only female, 9-month Tg-SwDI mice had overall reduced GSK3β phosphorylation compared to controls across all times of day (main effect of Genotype, p < 0.05). These data suggest that aged, female Tg-SwDI mice have increased levels of active GSK3β.

### Day/night differences in synaptic and intrinsic properties of hippocampal CA1 pyramidal cells are lost in Tg-SwDI mice

Brain circuits crucial for learning and memory reside in the hippocampus, which is one of the earliest regions impacted by AD (Braak and Braak, 1995; Mosconi, 2005; Pini et al., 2016). Post-mortem tissue analysis from patients with AD shows reduction of interneuron populations in the hippocampus (Brady and Mufson, 1997; Chan-Palay, 1987), and mouse models of AD have demonstrated the importance of interneuron function in cognitive performance (Palop and Mucke, 2016). Parvalbumin-expressing interneurons, in particular, are necessary for spatial working memory (Murray et al., 2011). Studies in multiple AD mouse models suggest that deficits in synaptic transmission occur early in the hippocampus (Hsia et al., 1999; Moechars et al., 1999; Palop et al., 2007). Given that hippocampal neurophysiology also varies across time-of-day (for review, see (Paul et al., 2020)), we hypothesized that diurnal hippocampal neuronal physiology may be disrupted in the Tg-SwDI mouse model in a manner that results in decreased inhibition.

To test this hypothesis, we recorded pharmacologically isolated spontaneous inhibitory postsynaptic currents (sIPSCs) using whole-cell voltage clamp of CA1 pyramidal cells (PCs) from 4-month-old male and female WT controls and Tg-SwDI mice (Figure 4A). In slices from WT control mice, sIPSC interevent interval (IEI) was significantly shorter during the day than at night (p < 0.001; Figure 4B (top)), and amplitude was significantly greater during the day than at night (p < 0.001; Figure 4B (bottom)). Because IEI is the inverse of frequency, this result indicates that the inhibition onto CA1 PCs was greater during the day than at night in controls, with more frequent and larger IPSCs. In recordings from CA1 PCs of Tg-SwDI mice, sIPSC amplitude was also greater during the day than at night (p < 0.001; Figure 4B (bottom)), similar to control mice. However, the day/night difference in IEI was reversed in slices from Tg-SwDI mice, with increased IEI during the day (p < 0.001; Figure 4B (top)), suggesting that TgSwDI mice have reduced inhibitory input onto excitatory PCs during the day.

**Figure 4.**
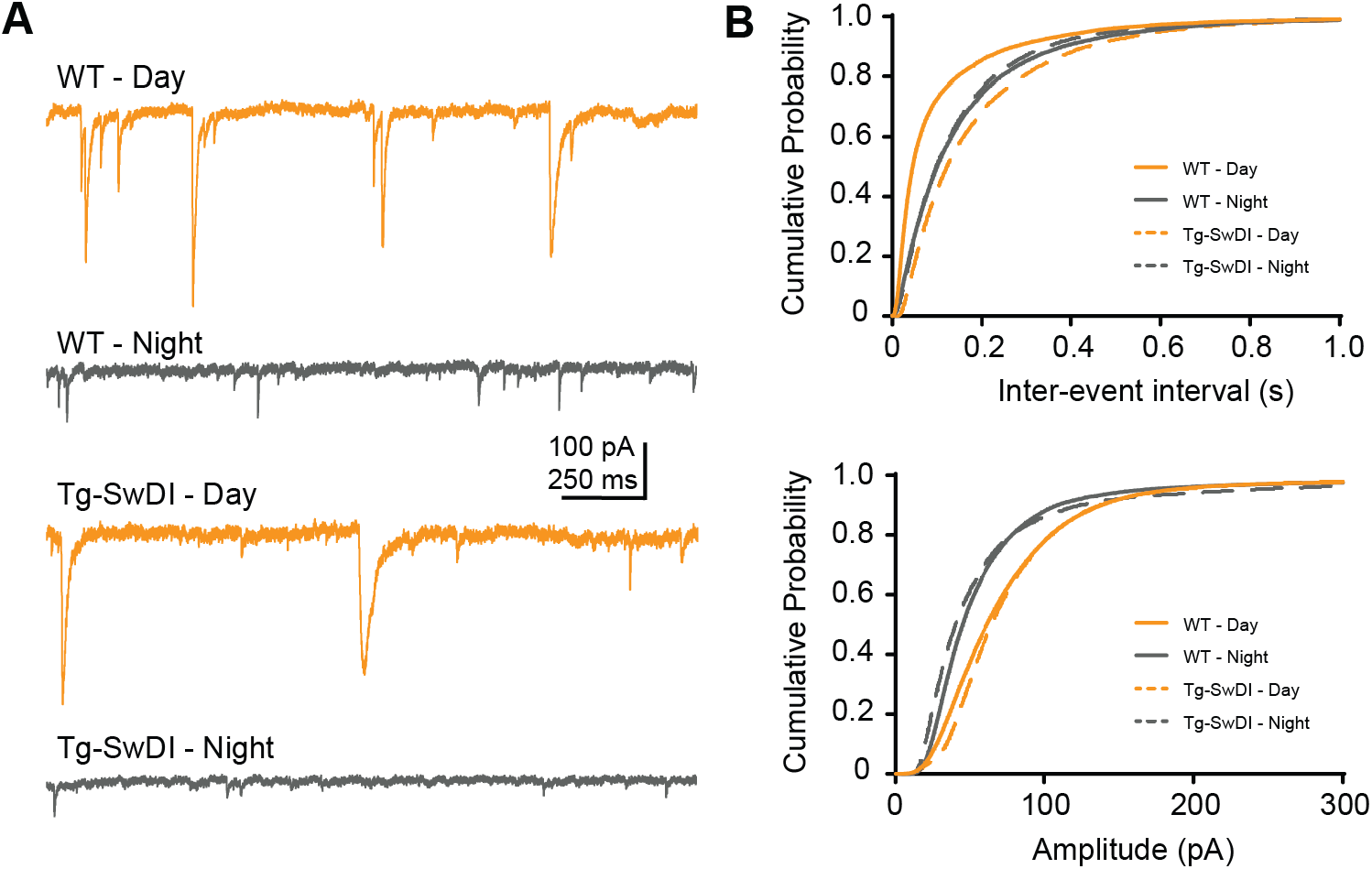
Increased daytime inhibitory input to CA1 pyramidal neurons impaired in Tg-SwDI mice. Whole cell patch clamp was used to measure amplitude and interevent interval (IEI) of sIPSCs in CA1 PCs after isolation with NBQX and CPP. Pipette recording solution contained CsCl and QX-314, and cells were held at −70mV. Representative traces (A) from WT (top) and Tg-SwDI (bottom) PCs at day (orange) and night (grey). Day/night differences in sIPSC inter-event interval (B, top) is reversed in Tg-SwDI mice. Day/night difference in amplitude (B, bottom) is preserved in Tg-SwDI mice (K-S test, Bonferroni-corrected alpha at p < 0.0125, n = 7-10 cells/group).

Next, we examined day/night differences in the excitability of CA1 PCs. Similar to data previously published in rats (Naseri Kouzehgarani et al., 2019), whole-cell current clamp recordings in 4-month-old control mice indicated a significant day/night difference in RMP, with greater depolarization at night (−66.23 ± 0.99 mV) compared to day (−70.00 ± 1.22 mV; t_51_ = −2.38, p < 0.05). To determine whether there were additional day/night differences in excitability, we assessed the number of action potentials generated in response to current injections of increasing magnitude from resting membrane potential, in the presence of vehicle (0.002% DMSO) (Figure 5A, right) or GABA receptor (GABAR) antagonists (Figure 5A, left) (10µM bicuculline and 1µM CGP55848). Three-way ANOVA of vehicle-treated slices indicated a significant current Step X Treatment X Time of Day interaction (p < 0.001, Supplementary Table 4). In the vehicle-treated group, CA1 PCs produced more action potentials at night than during the day in response to increasing current injections (for Steps 160 to 280 pA, p < 0.05; Figure 5A, left) and were thus more excitable at night.

**Figure 5.**
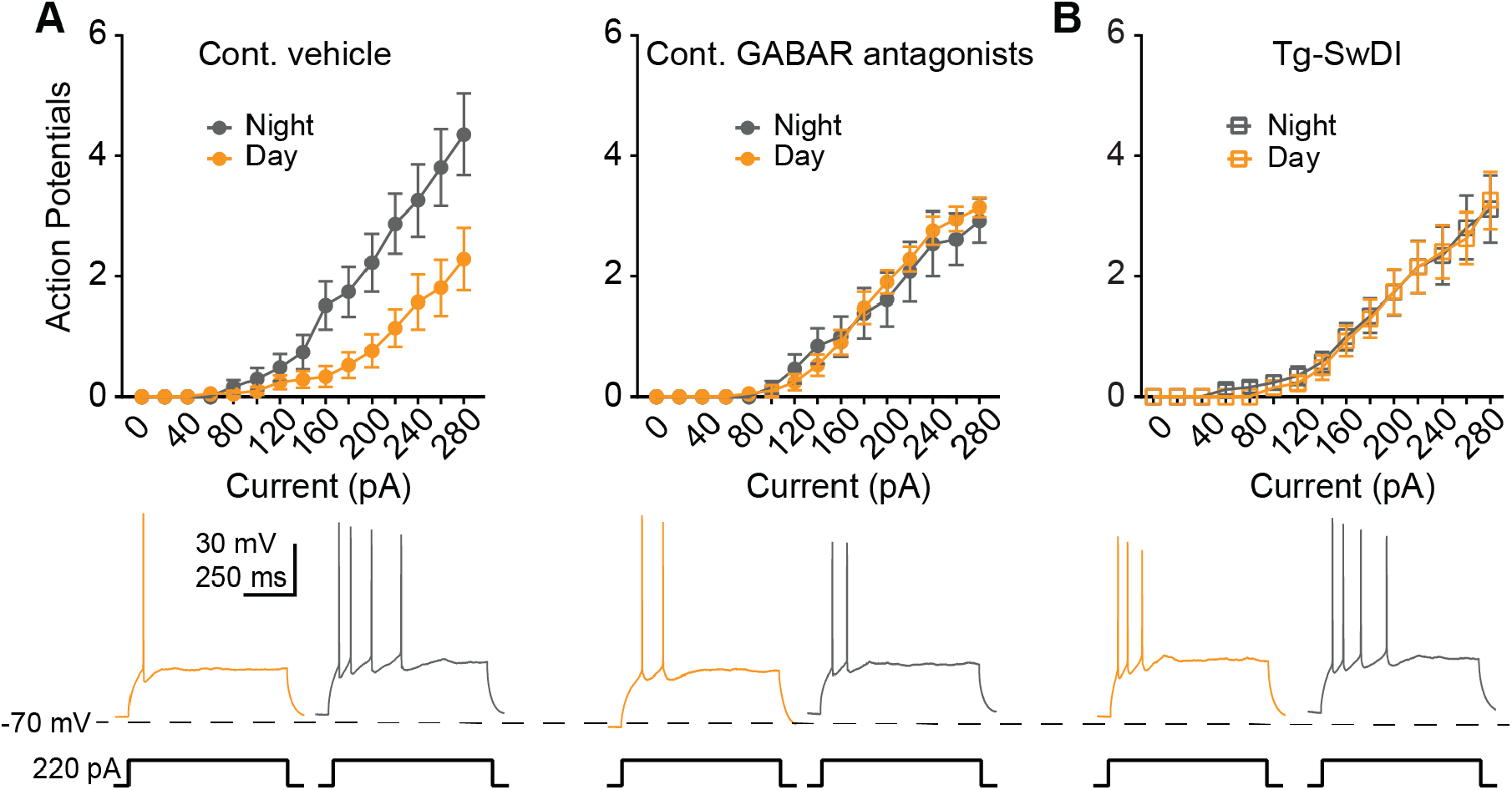
Loss of day/night differences in induced spike rate of hippocampal CA1 pyramidal cells in Tg-SwDI mice. Whole cell patch clamp was used to measure spikes per step with increasing current injections in CA1 PCs. Pipette recording solution contained KGlu, and cell voltage was not clamped. A. Whole-cell current clamp recordings of the number of spikes per step when recorded during the Day or Night in Per1::GFP mice in the absence (left; n = 21 cells (Day, orange), 31 cells (Night, grey)) or presence (right; n = 21 cells (Day), 13 cells (Night)) of GABA receptor antagonists; representative traces below (day: orange; night: grey). Three-way ANOVA indicated a significant Current Step X Treatment X Time of Day interaction (p < 0.001). In the vehicle-treated group, CA1 PCs produced more action potentials at Night compared to Day in response to increasing current injections (for Steps 160-to 280 pA, p < 0.05). In the presence of GABAR antagonists, the day/night difference in excitability was lost (simple main effects of Day vs Night at all current steps, p > 0.435). B. Pyramidal cell recordings from Tg-SwDI mice show loss of a day/night difference in induced spikes per step (p = 0.844; n = 27 cells (Day), 26 cells (Night)); representative traces below. See Supplementary Table 4 for statistical results.

In these vehicle-treated slices, both excitation and inhibition were present, constituting an intact local circuit. Therefore, to determine the role of inhibition in these day/night excitability differences, we also performed this experiment in the presence of GABAR antagonists. Blocking GABARs eliminated the day/night difference in CA1 PC excitability (simple main effects of day vs night at all current steps, p > 0.435; Figure 5A, right). Simple main effects comparing GABAR antagonist-treated cells to vehicle-treated cells during the day indicated statistical trends for the GABAR antagonist group to have more spikes at 180, 200, and 220 pA steps (p = 0.071, 0.055, and 0.067, respectively). Blocking GABARs had no effect on spikes per step at night.

We next asked if the day/night difference observed in vehicle-treated PCs was diminished in Tg-SwDI mice. There was no day/night difference in spikes per step in CA1 PCs in Tg-SwDI mice (p > 0.105; Figure 5B). Taken together, these data indicate that there is normally a day/night difference in CA1 PC excitability that requires inhibitory GABAergic transmission, as it is blocked by GABA receptor antagonists. Interestingly, the effect of blocking GABA receptors was very similar to the excitability phenotype observed in Tg-SwDI mice, with loss of the normal day/night difference in excitability in both conditions.

Finally, we probed for markers of interneurons, including GAD-67 and parvalbumin (PV), in the hippocampus of 4-month WT and Tg-SwDI mice. There was no significant difference in GAD^+^ fluorescence (t_11_ = 1.671, p > 0.05; Figure 6A-B) or in PV^+^ cell numbers (t_40_ = 1.135, p > 0.05; Figure 6C-D) between WT and Tg-SwDI mice.

**Figure 6.**
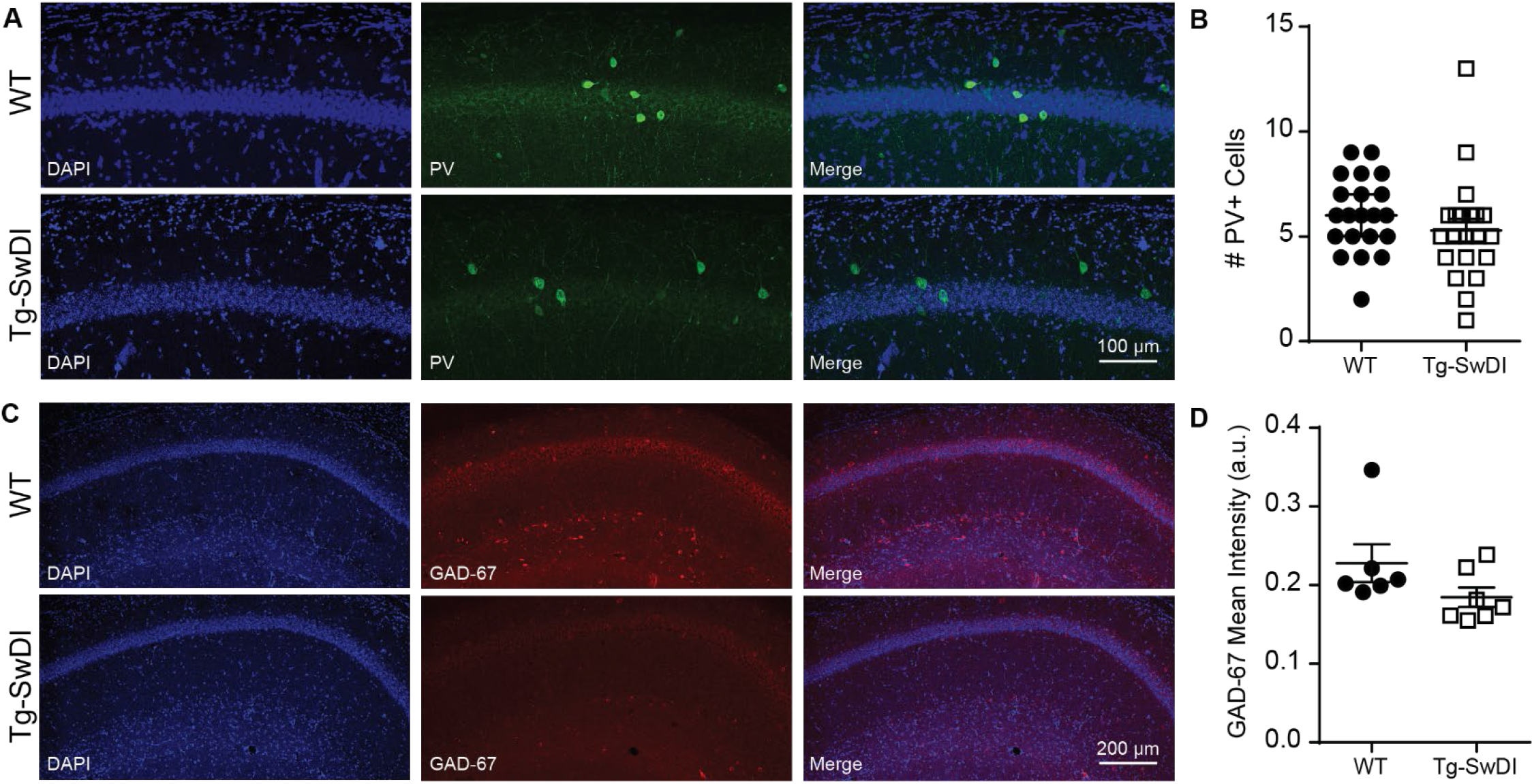
No change in interneuron markers in area CA1 of Tg-SwDI mice. Immunohistochemistry for GAD-67 and Parvalbumin from ZT 3-6 in CA1 of Tg-SwDI and WT mice. A. DAPI (left) and GAD-67 (middle) fluorescence in a coronal hippocampal slice from WT (top) and Tg-SwDI (bottom) mice and merged (right). Scale bar = 200 µm. B. Quantification of mean intensity, p > 0.05, n = 21 sections/group. C. DAPI (right) and parvalbumin (middle) fluorescence in a coronal slice from WT (top) and Tg-SwDI (bottom) mice merged (right). Scale bar = 100 µm. D. Quantification of PV^+^ cell count, p > 0.05, (n = 21 sections/genotype).Biocytin filled CA1 PCs from WT (left) and Tg-SwDI (right) mice.

## DISCUSSION

Altered rest and activity patterns are commonly reported symptoms in patients with AD (Ancoli-Israel et al., 1997; Harper et al., 2004; Harper et al., 2005; Mishima et al., 1997; Mishima et al., 1999; Musiek et al., 2018; Okawa et al., 1991; Satlin et al., 1995; Vitiello and Prinz, 1989; Witting et al., 1990). This circadian disruption may exacerbate AD pathology and vice versa (Musiek, 2017). In the APP-PS1 mouse model of AD, adult-specific genetic ablation of the molecular clock accelerates amyloid beta pathology (Kress et al., 2018). Sleep deprivation, in both wild type and AD models, potentiates amyloid plaque formation (Kang et al., 2009; Roh et al., 2012). In the Tg-SwDI model, utilized in this paper, circadian locomotor activity is disrupted and day/night differences in excitability of SCN neurons are diminished due to reduced daytime firing (Paul et al., 2018). Additionally, cerebral microvascular amyloid is first detected in the hippocampus, subiculum, and cortex at 3 months in Tg-SwDI mice with pronounced spread by 6 months, covering the entire forebrain by 12 months (Davis et al., 2004; Xu et al., 2014). In the present study, we show lack of the normal diurnal differences in spatial working memory in this AD model, which develop relatively early in parallel with a general reduction in hippocampal molecular clock gene expression. Importantly, we also provide the first evidence that normal day/night differences in hippocampal neurophysiology, including inhibitory synaptic input, are impaired in a mouse model of AD.

Given the reported circadian behavior and SCN disturbances in the Tg-SwDI model, we aimed to investigate whether this model also had disrupted rhythms in spatial working memory. While the Tg-SwDI model has been shown to develop deficits in Barnes maze performance beginning at 3 months of age (Xu et al., 2007; Xu et al., 2014), there had been no published studies of day/night differences in performance of any tasks at any age. Additionally, the Barnes maze requires multiple daily trials over the course of multiple days (Pitts, 2018), introducing the possible confounds of light exposure or sleep deprivation. Therefore we chose to assess spatial working memory with T-maze Spontaneous Alternation, a task that can be measured at two different times of day with minimal training and normally shows strong day/night difference in performance (Davis et al., 2020; Ruby et al., 2013; Ruby et al., 2008). As previously shown, we observed a significant day/night difference in performance in controls, with high alternation at night. However, female Tg-SwDI animals had lower alternation than controls at night with no significant difference between day and night performance. At 9 months, both male and female wild-type mice had significantly higher performance at night than during the day, while the Tg-SwDI mice had no significant day/night difference. In healthy individuals, performance in hippocampal-dependent tasks is best after waking and declines after several hours (Wright et al., 2006). Given the significant circadian disruption seen in AD patients, daily patterns of cognition are an important consideration for treatment of AD. Approximately 25% of patients with AD exhibit sundowning, a clinical phenomenon in which neuropsychiatric symptoms manifest in the late afternoon, evening or at night (Khachiyants et al., 2011). Additionally, patients in the preclinical stage of Alzheimer’s disease are more likely to have increased rest/activity rhythm fragmentation (Musiek et al., 2018). Exploring circadian deficits present in AD underlines a growing need to design therapies for AD that could be administered at specific times of day for optimal benefit.

Because the day/night difference in spatial working memory is linked to the molecular clock (Shimizu et al., 2016; Snider et al., 2016; Snider and Obrietan, 2018), we hypothesized that expression of circadian clock genes and clock-controlled genes may be altered in AD. The molecular clock consists of several core clock proteins, including positive regulators CLOCK and BMAL1, which dimerize in the nucleus and bind to E-box regions of clock genes *Period (Per)* and *Cryptochrome (Cry)*, the negative regulators, allowing their transcription. The transcripts are then translocated to the cytoplasm where they are translated into proteins. While in the cytoplasm, PER and CRY accumulate and are either marked for degradation or re-entry to the nucleus by phosphorylation via casein kinase 1 or GSK3β, with these phosphorylation events setting the period length (Besing et al., 2015; Lee et al., 2011). Eventually, PER and CRY dimerize and are shuttled back to the nucleus where they interfere with the binding of CLOCK and BMAL1, thus turning off their own transcription. This entire process takes ∼24 hours. A second, auxiliary loop exists with the activator retinoid-related orphan receptor (ROR) and repressor REV-ERBαβ, which drives the transcription of *Bmal1* (Partch et al., 2014; Rosensweig and Green, 2020). In addition to these core clock components, the molecular clock also drives rhythmic expression of clock-controlled genes in a region-specific manner throughout the brain, including the hippocampus (Paul et al., 2020; Renaud et al., 2015).

Reductions in several core clock components as measured by both mRNA and protein have been reported in multiple AD models. Our findings are consistent with other studies showing reduced clock gene expression and protein levels in both cell culture and mouse models. Two other AD mouse models (APP/PS1 and 5XFAD) showed reduced *Per2* mRNA and/or protein levels in the SCN or hypothalamus, compared to age-matched controls (Duncan et al., 2012; Song et al., 2015). In the hippocampus, intracranial injection of Aβ_31-35_ in WT mice reduced the nocturnal peak in PER2 protein levels (Wang et al., 2016). In young APP/PS1 mice, hippocampal clock gene expression was not different from WT controls, with both genotypes showing a time of day difference (Oyegbami et al., 2017). However, by 12-15 months, APP/PS1 mice showed reduced diurnal rhythms in hippocampal clock gene mRNA expression (Ma et al., 2016). Our present study shows overall reduction in gene expression of *Bmal1, Bmal2, Cry1, Per1*, and *Reverbα*, across both males and females, and reduced *Per2* expression specific to female mice at 4 months of age. Thus, expression of many but not all molecular clock genes and protein levels (PER2) were lower overall in the Tg-SwDI hippocampus than in controls (with the exception of *Cry2* mRNA and BMAL1 protein levels). We do not yet understand the downstream implications of reduced clock gene expression in AD mouse models.

In addition to reduced expression of core components of the molecular clock, we observed reduced levels of phosphorylated (inactive) GSK3β, which phosphorylates various components of the molecular clock (Besing et al., 2015; Iitaka et al., 2005; Iwahana et al., 2004), in older female Tg-SwDI mice. Younger mice (both Tg-SwDI and WT) and older males had normal diurnal variation of pGSK3β in the hippocampus as previously reported (Besing et al., 2017). Because the GSK3 de-phosphorylated state is the active state, reduced phosphorylation levels would suggest that the AD females have increased activity of GSK3β in the hippocampus. This activity can lead to increased Aβ and tau in the brain and alter protein homeostasis (Leng et al., 2019). However, reduced pGSK3β was not observed in 4 month old Tg-SwDI mice, suggesting that the early deficits observed here and by others (Aβ accumulation, cognitive deficits, and reduced molecular clock components) are not due to altered GSK3β rhythms, and there must be another currently unidentified mechanism at this age. Some evidence suggests reduced clock function may lead to several issues including Aβ accumulation, synaptic dysfunction, oxidative stress, altered proteostasis, immune dysfunction, and neuroinflammation (Leng et al., 2019). Interestingly, neuroinflammation may be a mediator of early cognitive dysfunction in Tg-SwDI such that microvascular amyloid in hippocampal-associated regions (i.e., the subiculum) may lead to hippocampal neuroinflammation and impaired function (Robison et al., 2019; Xu et al., 2007).

Importantly, the molecular clock exerts transcriptional regulation of excitable membranes. Many ion channel-related genes examined in CircaDB database show significant 24-hour rhythms in expression in the SCN (Pizarro et al., 2013). Therefore, it is possible that deficits in the molecular clock alter neuronal membrane physiology. A substantial literature illustrates the existence of daily rhythms in neuronal physiology in multiple regions of the rodent brain (for review, see (Paul et al., 2020)). The ratio of excitation to inhibition is crucial for normal brain function and modelling based on human EEG data suggests that this ratio exhibits circadian differences (Chellappa et al., 2016). Notably, the ratio of excitation to inhibition is disrupted early in AD models (Palop et al., 2007; Roberson et al., 2011). In the J20 mouse model of AD, daytime recordings of sIPSCs onto PCs of the parietal cortex showed a reduction in frequency compared to NT littermate controls (Verret et al., 2012). When inhibitory PV interneuron excitability in the J20 model was restored, sIPSC amplitude and frequency was similar to NT controls, and epileptiform activity was reduced (Verret et al., 2012). Our study is the first to report the lack of day/night differences in sIPSCs in a mouse model of AD.

Circadian rhythms in neurophysiology have been best described in the SCN, where the action potential firing rate of neurons is higher during the day compared to night (Colwell, 2011; Paul et al., 2016). In Tg-SwDI mice, the daytime firing rate of SCN neurons is reduced, resulting in a dampening of the day/night difference (Paul et al., 2018). Thus far, a single report has shown that rat hippocampal neurons exhibit day/night differences in one measure of excitability, with CA1 pyramidal cell RMP being more depolarized during the night compared to day (Naseri Kouzehgarani et al., 2019). In control mice, we demonstrated a similar day/night difference in excitability, with CA1 pyramidal cells resting at more hyperpolarized potentials during the day. Additionally, we showed that current injections of increasing magnitude into the soma of CA1 pyramidal cells produce more action potentials during the night, suggesting an overall increase in CA1 PC excitability at night. Importantly, we showed for the first time an absence of day/night difference in excitability in an AD mouse model, suggesting a circadian dysregulation of excitability. In future studies, it will be important to investigate if the lack of the day/night difference in CA1 PC excitability that we report here is tied to the dysregulation of the molecular clock. Notably, in control slices, blocking GABAR signaling abolished the day/night difference in CA1 PC excitability. Altogether, our data suggest that both cell autonomous (potential molecular clock-mediated rhythms in ion channel function) and non-autonomous (circadian regulation of tonic or phasic inhibition) mechanisms may work in concert to maintain temporal control of excitability, which is lost in pathophysiological conditions such as AD.

The lack of day/night differences in CA1 PC excitability and sIPSCs in TgSWDI mice align with several pieces of evidence supporting the hypothesis of hyperexcitability in AD. Epileptiform activity has been documented in AD patients and is more likely to be observed at night, when patients are asleep (Anderson et al., 2015; Vossel et al., 2016). Several populations of interneurons, including parvalbumin-expressing interneurons, are reduced in post-mortem tissue of AD patients (Brady and Mufson, 1997; Chan-Palay, 1987). Fast-spiking parvalbumin interneurons are necessary for generation of oscillations in the gamma frequency, which are important in learning and memory (Cardin et al., 2009; Nyhus and Curran, 2010; Sohal et al., 2009; Stam et al., 2002). Deficits in gamma oscillatory activity and reduced activity of fast-spiking interneurons are observed in both mouse models of AD and AD patients (Cheng et al., 2020; Martinez-Losa et al., 2018; Verret et al., 2012; Vossel et al., 2016). Thus, future work should investigate circadian dysregulation of specific interneuron sub-types in the context of AD.

Our present findings highlight the importance of circadian regulation of neuronal physiology and the potential for chronotherapeutic interventions. Clarifying how AD progression impacts day/night neurophysiological differences could lead to chronotherapeutic strategies, whereby hyperexcitability is treated at a specific time of day. In the future, it will be important to dissect roles of somatic-targeting and dendritic-targeting interneurons to determine where the loss of inhibition is originating. Because CSF Aβ and tau exhibit a diurnal pattern (Lucey et al., 2015; Roh et al., 2012), other AD biomarkers are also likely to differ in the day versus night. Thus, the circadian clock is an important consideration for all future studies of AD pathogenesis and therapeutics.

## ACKNOWLEDGEMENTS

This work was supported by NIH grants R01NS082413 (KLG), R56AG061785 (EDR and KLG), RF1AG059405 (EDR), R01NS108713 (KLG and RMC), T32NS061788 (ARF), and F31NS115299 (LKG). We would like to thank Allie Widman for technical assistance with electrophysiology and Courtney Rogers for assistance with animal behavior experiments. We would also like to thank Kavitha Abiraman for helpful comments in the writing of the manuscript.

---

**Supplementary Figure 1.**
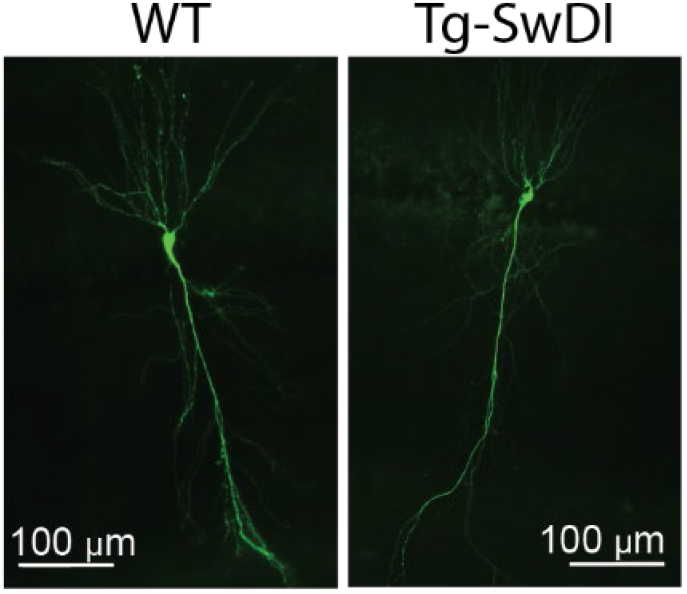
Biocytin filled CA1 PCs from WT (left) and Tg-SwDI (right) mice.

**Supplementary Figure 2.**
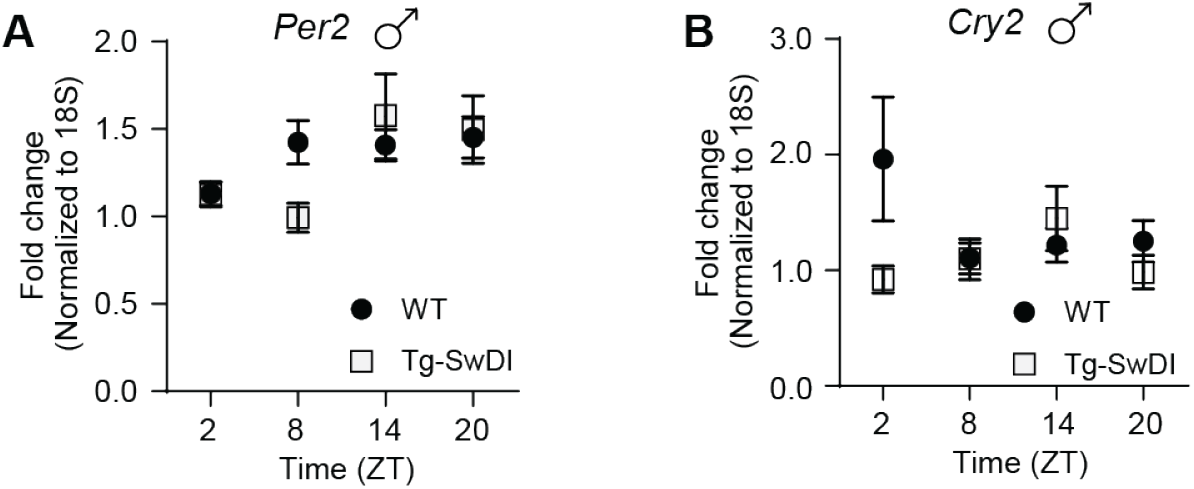
*Per2* mRNA expression in 4-month male mice. Transcript levels from WT (filled circles) and Tg-SwDI (open squares) male mice at ZT 2, 8, 14, and 20 for *Per2* (A) and *Cry2* (B). Two-way ANOVA indicated no significant effects. See Supplementary Table 2 for statistical results.

**Supplementary Figure 3.**
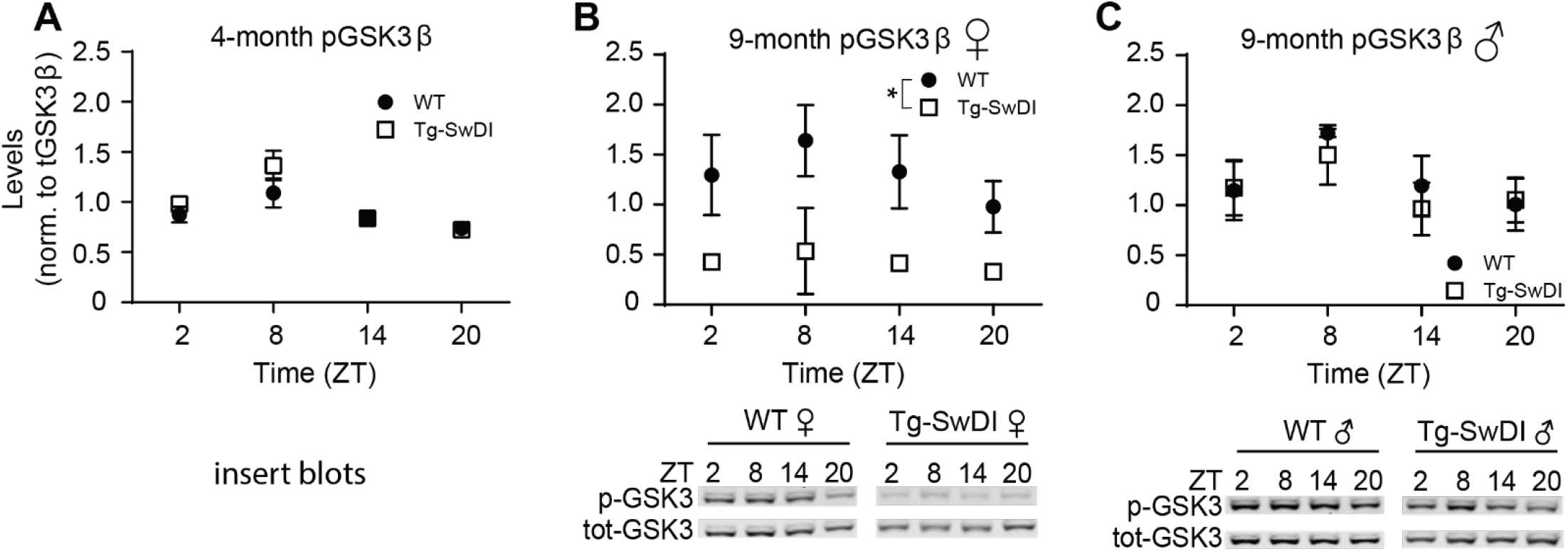
Loss of day/night variation pGSK3β expression in Tg-SwDI mice. Normalized expression for pGSK3β in Tg-SwDI mice (open squares) and age-matched WT controls (solid circles). For levels of p-GSK3β (normalized to total GSK3β), ZT 8 levels were greater than all other time points (significant main effect of Time-of-Day and post hoc, p < 0.01), with no effect of Genotype or significant interaction (n = 8-15/group). At 9-months females (B) showed a significant effect of Genotype (p < 0.05; no other effects) but no genotype difference was detected in male (C) mice (n = 5-9 mice/group). See Supplementary Table 3 for statistical results.

**Table 1.**

|  | Genotype | Time of Day | Interaction |
| --- | --- | --- | --- |
| 4-month females | $\chi^2 (1) = 3.117, p = 0.077$ | $\chi^2 (1) = 8.857, p = 0.003$ | $\chi^2 (1) = 12.076, p = 0.001$ |
| 4-month males | $\chi^2 (1) = 0.193, p = 0.660$ | $\chi^2 (1) = 6.003, p = 0.014$ | $\chi^2 (1) = 1.633, p = 0.201$ |
| 9-month females | $\chi^2 (1) = 0.663, p = 0.415$ | $\chi^2 (1) = 1.030, p = 0.310$ | $\chi^2 (1) = 6.852, p = 0.009$ |
| 9-month males | $\chi^2 (1) = 0.268, p = 0.605$ | $\chi^2 (1) = 3.136, p = 0.077$ | $\chi^2 (1) = 8.526, p = 0.004$ |

**Table 2.**

|  | Genotype | Time of Day | Interaction |
| --- | --- | --- | --- |
| Bmal1 | $F_{1, 104} = 12.522, p = 0.001$ | $F_{3, 104} = 1.695, p = 0.173$ | $F_{3, 104} = 1.071, p = 0.365$ |
| Bmal2 | $F_{1, 104} = 16.301, p < 0.001$ | $F_{3, 104} = 0.781, p = 0.507$ | $F_{3, 104} = 0.478, p = 0.698$ |
| Cry1 | $F_{1, 104} = 24.517, p < 0.001$ | $F_{3, 104} = 2.231, p = 0.089$ | $F_{3, 104} = 0.165, p = 0.919$ |
| Cry2 females | $F_{1, 45} = 6.844, p = 0.012$ | $F_{3, 45} = 0.583, p = 0.629$ | $F_{3, 45} = 1.107, p = 0.356$ |
| Cry2 males | $F_{1, 49} = 1.641, p = 0.206$ | $F_{3, 49} = 0.657, p = 0.582$ | $F_{3, 49} = 1.957, p = 0.133$ |
| Per1 | $F_{1, 104} = 19.791, p < 0.001$ | $F_{3, 104} = 0.460, p = 0.711$ | $F_{3, 104} = 0.650, p = 0.585$ |
| Per2 females | $F_{1, 45} = 19.938, p < 0.001$ | $F_{3, 45} = 8.189, p < 0.001$ | $F_{3, 45} = 0.974, p = 0.414$ |
| Per2 males | $F_{1, 51} = 0.368, p = 0.547$ | $F_{3, 51} = 4.670, p = 0.006$ | $F_{3, 51} = 2.209, p = 0.098$ |
| Rev-erb alpha females | $F_{1, 45} = 21.708, p < 0.001$ | $F_{3, 45} = 0.537, p = 0.659$ | $F_{3, 45} = 0.305, p = 0.822$ |
| Rev-erb alpha males | $F_{1, 51} = 7.104, p = 0.010$ | $F_{3, 51} = 0.383, p = 0.766$ | $F_{3, 51} = 1.067, p = 0.371$ |
| 18S females | $F_{1, 45} = 6.568, p = 0.327$ | $F_{3, 45} = 0.779, p = 0.512$ | $F_{3, 45} = 0.395, p = 0.757$ |
| 18S males | $F_{1, 51} = 0.978, p = 0.327$ | $F_{3, 51} = 0.057, p = 0.982$ | $F_{3, 51} = 0.748, p = 0.529$ |

**Table 3.**

| 4 months | Genotype | Time of Day | Interaction |
| --- | --- | --- | --- |
| BMAL1 | $F_{1,100} = 0.0065, p = 0.940$ | $F_{3,100} = 18.602, p < 0.001$ | $F_{3,100} = 5.214, p = 0.002$ |
| PER2 | $F_{1,98} = 9.357, p = 0.003$ | $F_{3,98} = 3.913, p = 0.011$ | $F_{3,98} = 0.871, p = 0.459$ |
| pGSK3-β norm to tGSK3-β | $F_{1,89} = 2.242, p = 0.138$ | $F_{3,89} = 11.584, p < 0.001$ | $F_{3,89} = 1.160, p = 0.330$ |
| 9 months |  |  |  |
| BMAL1 | $F_{1,95} = 6.553, p = 0.012$ | $F_{3,95} = 2.249, p = 0.088$ | $F_{3,95} = 0.754, p = 0.523$ |
| PER2 | $F_{1,95} = 9.764, p = 0.002$ | $F_{3,95} = 2.331, p = 0.079$ | $F_{3,95} = 0.073, p = 0.974$ |
| pGSK3-β females | $F_{1,44} = 25.568, p < 0.001$ | $F_{3,44} = 0.994, p = 0.404$ | $F_{3,44} = 0.274, p = 0.844$ |
| pGSK3-β males | $F_{1,44} = 0.191, p = 0.664$ | $F_{3,44} = 1.646, p = 0.193$ | $F_{3,44} = 0.125, p = 0.945$ |

**Table 4.**

|  |  |
| --- | --- |
|  | WT Vehicle vs GABA blockers |
| Current Step | $F_{14, 1148} = 75.038, p < 0.001$ |
| Current Step X Treatment | $F_{14, 1148} = 0.352, p = 0.987$ |
| current step X Time of Day | $F_{14, 1148} = 1.775, p = 0.038$ |
| Current Step X Treatment X Time of Day | $F_{14, 1148} = 3.320, p < 0.001$ |
| Treatment | $F_{1, 82} = 0.037, p = 0.848$ |
| Time of Day | $F_{1, 82} = 2.711, p = 0.103$ |
| Treatment X Time of Day | $F_{1, 82} = 3.425, p = 0.068$ |
|  | Tg-SwDI |
| Current Step | $F_{14, 714} = 46.351, p < 0.001$ |
| current step X Time of Day | $F_{14, 714} = 0.067, p = 1.000$ |
| Time of Day | $F_{1, 51} = 0.29, p = 0.864$ |

